# Consequences of dietary amino acid imbalance on growth and gas exchange in black soldier fly (*Hermetia illucens*) larvae

**DOI:** 10.1101/2025.09.05.674443

**Authors:** M. Schow-Madsen, K. Jensen, M. Márquez-Toribio, M.L. Schøn, T.M. Schou, J.V. Nørgaard, J. Overgaard

## Abstract

Black soldier fly larvae (*Hermetia illucens* (L.); BSFL) production relies on efficient nutrient conversion into protein-rich biomass to ensure sustainability and economic viability. While the effects of dietary protein content and macronutrient ratios are well described, the metabolic costs of providing an imbalanced essential amino acid (EAA) supply in dietary proteins remain poorly understood. We examined how dietary EAA imbalance affects growth, metabolism, and energy allocation in BSFL. Larvae were reared on a reference diet (chicken feed) and three isonitrogenous, semi-synthetic diets: one balanced diet matching the EAA profile of the reference, and two unbalanced diets selectively deficient in either leucine, phenylalanine, and threonine or isoleucine, lysine, and tryptophan. Using open flow respirometry, daily growth monitoring, and biochemical analysis, we quantified gas exchange, body composition, and the energetic costs of biomass and protein deposition. EAA imbalance delayed development, reduced growth and protein content, and promoted lipid accumulation, while O_2_ consumption and CO_2_ production were reduced and respiratory exchange ratios exceeded 1.0, indicating elevated lipogenesis. Metabolic costs per unit protein deposited were higher on EAA-deficient diets (up to 3.63 kJ per kJ protein), reflecting inefficient amino acid utilisation. In contrast, the balanced diet supported more efficient protein deposition at a lower energetic cost (2.73 kJ per kJ protein). Our findings demonstrate that suboptimal dietary EAA profiles reduce metabolic efficiency in BSFL. Quantifying the physiological costs of amino acid imbalance provides benchmarks to guide feed formulations in insect production systems to minimise input costs and maximise the nutritional quality and quantity of larval output.

## Introduction

Optimising insect nutrition is crucial in insects produced for food and feed to maximize feed conversion and utilization efficiency to minimize production costs (Oonincx *et al*., 2025). Black soldier fly larvae (*Hermetia illucens* (L.); BSFL) are increasingly applied to support circular waste management as they convert low-value organic side streams into protein- and lipid-rich biomass for animal feed (Liu *et al*., 2019; Purkayastha and Sarkar, 2022; van der Heide *et al*., 2021) and produce organic plant fertiliser from the frass (Lomonaco *et al*., 2024). Rapid growth, high feed conversion efficiency, and tolerance to a wide range of low-quality substrates make BSFL well-suited for large-scale production systems (Liu *et al*., 2019; Siddiqui *et al*., 2024). However, production efficiency - including biomass conversion rate, larval yield, growth rate, survival, and body composition - is strongly modulated by the nutritional quality of the feed (Gold *et al*., 2018; Liu *et al*., 2019). Research on BSFL has explored diverse feedstocks and underutilised organic side streams (Broeckx *et al*., 2021; Gold *et al*., 2018; Oonincx *et al*., 2015), and substantial progress has been made in understanding the effects of dietary protein and carbohydrate content (Barragan-Fonseca *et al*., 2019; Cammack and Tomberlin, 2017; Schøn *et al*., 2025a). As BSFL production scales up, it is increasingly important to evaluate how suboptimal diets affect growth, metabolism, and body composition, to develop cost-effective feeds that optimise larval yield and nutritional value while maintaining economic viability.

Despite increasing attention to BSFL nutrition, the role of micronutrient balance, especially essential amino acids (EAAs), remains insufficiently understood (Tomberlin *et al*., 2023). EAAs are required for protein synthesis and cannot be synthesised *de novo* (Wu *et al*., 2014). In BSFL, few studies have tested individual EAA limitations, showing that deficiencies or imbalances constrain somatic growth and delay development (Berggreen *et al*., 2025; Koethe *et al*., 2022; Spranghers *et al*., 2025). However, the broader metabolic consequences of imbalanced dietary EAA profiles, including effects on energy allocation and protein assimilation, remain unexplored. When EAAs are limiting, excess non-limiting amino acids cannot support protein synthesis and are instead deaminated with a large proportion of the resulting carbon redirected into lipid synthesis (Berggreen *et al*., 2025; Stryer, 1995), potentially increasing the metabolic cost of growth.

Growth is metabolically costly, requiring energy for biomass synthesis, tissue maintenance, and cellular turnover (Eriksen, 2022; Laganaro *et al*., 2021; Schmidt-Nilsen, 1997). It is well-described that the energetic cost of digesting and assimilating protein exceeds that of carbohydrates and lipids (Goodrich *et al*., 2024; Hawkins, 1991; Secor, 2009), and when protein is present but poorly utilisable, a greater proportion of total energy metabolism is allocated toward maintenance rather than growth (Laganaro *et al*., 2021). As feed accounts for a major part of the BSFL production costs, efficient conversion of diverse, low-value substrates into high-protein biomass is key to economic viability. High-quality substrates accelerate growth and improve larval protein content and nutrient composition (Barragan-Fonseca *et al*., 2019; Schøn *et al*., 2025a), increasing their nutritional value. In contrast, suboptimal feeds prolong development to pupation and reduce both biomass yield and quality (Barragan-Fonseca *et al*., 2019; Gold *et al*., 2018), though such diets may be cheaper and easier to formulate from low-cost side-stream. Whether the aim is to minimise feed input costs or to maximise the nutritional quality of the larval biomass, a clear understanding of the metabolic costs associated with nutrient assimilation is essential for evaluating feed efficiency. As BSFL are primarily valued for their protein content, reduced protein deposition directly lowers both the nutritional value of the larvae and the economic return. Identifying how suboptimal dietary EAA profiles influence the energetic cost of protein deposition is therefore important to guide feed formulation strategies in large-scale production.

The objectives of this study were to evaluate how amino acid balance affected BSFL growth, survival, and body composition, how respiratory activity changes during the last 7 days of production, and to estimate the energetic cost of lipid and protein deposition using oxygen-based calorimetry. This study evaluates the gross cost of growth, defined as the energy expended per unit of biomass gained, providing an integrative measure of metabolic efficiency during growth.

## Materials and methods

### Experimental animals and general design

*Hermetia illucens* larvae were provided by the commercial BSFL facility ENORM Biofactory A/S (Flemming, Denmark) and transported 1-hour to the laboratory facilities shortly before each experiment. The larvae used in the experiments were 5-6 days old with a mass of ~5 mg, corresponding to the 4^th^-5^th^ larval stage (Gligorescu *et al*. 2019).

The experiments were designed to examine how dietary deficiencies of EAAs affect the gross cost of growth in BSFL. To examine this, growth, gas exchange, body composition, and energy expenditure were compared in BSFL reared on four different diets (see below) for seven days at 27°C. Each replicate in each diet group contained 600 larvae, housed in 1 L respirometer chambers with ~340 g of feed. Over four weeks, measurements were performed using a blocked design, testing one or two replicates per diet weekly, resulting in seven replicates per diet. The experimental system recorded CO_2_ production and O_2_ consumption on an hourly basis, and each day, 15 larvae from each replicate were randomly selected, weighed and then returned to the chamber. At the end of the experiment, all larvae were weighed, counted, and stored at −20°C for later analysis of water, crude lipid, and crude protein content.

### Experimental diets

This experiment included a reference diet (REF), and three treatment diets - a balanced (B), and two unbalanced diets (UA and UB). The REF diet consisted solely of commercial chicken feed (KylleKræs 1, Danish Agro; Table S1), which contained amino acids, minerals, and vitamins to meet the nutrient requirements of growing chickens (23.0% protein, 65.4% carbohydrates, and 5.2% lipids). The REF diet was included to ensure that our rearing conditions supported growth and metabolism comparable to previous BSFL studies using high-productivity feeds (Schøn *et al*., 2025b). The three treatment diets (B, UA, UB) were formulated to explore the impact of EAA deficiencies (Table 1). The balanced diet (B) had the same amino acid composition as in chicken feed, but while chicken feed had a protein content of 20.3% of diet dry mass, the semi-synthetic diets had a protein plus free amino acid content of 15.2 % of diet dry mass to ensure a high requirement for balanced amino acids for growth (Berggreen *et al*., 2025). To create a highly imbalanced diet (UB) where we would be likely to measure metabolic effects of amino acid deficiency, we excluded tryptophan, isoleucine, and lysine in the synthetic part of the UB diet as these are the three EAAs that each limit growth the most when excluded from the EAA mix (Berggreen *et al*., 2025). To create a gradually less but still highly imbalanced diet (UA), we excluded leucine, threonine, and phenylalanine in the synthetic part of the UA diet, as these are the next individual EAAs with highest consistent growth limiting effect when deficient (Berggreen *et al*., 2025). All three treatment diets contained 35.1% chicken feed to ensure a base supply of essential macro- and micronutrients. Compared to the REF diet, the semi-synthetic treatment diets contained less total protein to promote high protein utilisation and induce a growth-limiting effect of EAAs, whose relative concentrations were below the levels required for maximal growth and protein deposition (Berggreen *et al*., 2025). Importantly, the B diet formulation has previously been validated for BSFL growth in Berggreen *et al*. (2025), confirming its suitability for controlled nutritional experiments. This chicken feed fraction in the treatment diets provided approximately 8.1% protein, 23% carbohydrates, and 1.8% lipids. The remaining 64.9% of the diet consisted of a carbohydrate mixture (sucrose and dextrin, 1:1), a free amino acid mixture, cellulose, and other components including Vanderzant vitamin mixture, Wesson’s salt mixture, linoleic acid (80% linoleic acid, 20% oleic acid), cholesterol, and nipagin (1:6:1:1:1). Cellulose served as a structural element, while nipagin functioned as a preservative.

**Table 1.** Ingredient and nutritional composition of the three semi-synthetic diets based on dry mass content. The three diets differed only in the specific free amino acid mix used.

|  | Diets |
| --- | --- |
| **Ingredients (%)** |  |
| Chicken feed<sup>1</sup> | 35.1 |
| Carbohydrate mix<sup>2</sup> | 39.3 |
| Amino acid mix<sup>3</sup> | 7.1 |
| Cellulose | 14.3 |
| Agar | 1.7 |
| Other additives<sup>4</sup> | 2.5 |
| **Nutrients (%)** |  |
| Carbohydrate | 78.3 |
| Digestible carbohydrates | 61.1 |
| Protein | 8.1 |
| Free amino acid mix | 7.1 |
| Lipids | 1.8 |
| Other components<sup>5</sup> | 4.7 |
| Energy (kJ/g)<sup>6</sup> | 16.3 |
| Usable energy (kJ/g)<sup>7</sup> | 13.6 |

<sup>1</sup>See Table S1.
<sup>2</sup>1:1 sucrose and dextrin.
<sup>3</sup>See Table 2.
<sup>4</sup>Other additives include Vanderzant vitamin mixture (0.25%), Wessons salt mixture (1.50%), nipagin (0.25%) and the lipids; linoleic acid (0.20%), oleic acid (0.05%) and cholesterol (0.25%).
<sup>5</sup>Micronutrients plus other additives.
<sup>6</sup>Energy is calculated using energy contents of 17 kJ/g for carbohydrates, 17 kJ/g for protein, and 37 kJ/g for lipids (Capuano *et al.*, 2018).
<sup>7</sup>Excludes the contribution from indigestible carbohydrates (fibre in chicken feed, cellulose, and agar).

In diet B, the amino acid profile was matched to that of REF, enabling a direct comparison of amino acids derived exclusively from chicken feed versus those provided as free crystalline amino acids. Diet UA was selectively deficient in leucine, phenylalanine, and threonine, whereas diet UB was deficient in isoleucine, lysine, and tryptophan (Table 2). These six amino acids each limited BSFL growth significantly when not included in the free amino acid mix (Berggreen et al. 2025), and their deficiencies were here combined in two random groups in the two diets. To maintain a constant amount of free amino acid of 7.1% of the dry diet (Table 1), non-essential amino acids were adjusted to compensate for the removed EAA. All diets were prepared by mixing 320 g of heated agar-water solution with 80 g of the designated dry ingredients. After solidification in the respiration chambers, some water evaporated, yielding 351.6 ± 5.5 g of wet diet (~77% water). Each chamber contained 600 BSFL, corresponding to ~1.71 larvae per gram of wet diet or 0.133 g of dry feed per larva.

**Table 2.** Calculated amounts of essential and non-essential amino acids per kg dry diet in the chicken feed (REF) diet and the three treatment diets varying in amino acid composition. The B diet contains all amino acids in similar proportions as in chicken feed (Table S1), but at lower concentration. The amino acids in bold stem only from the chicken feed part of the diet, as they were not added in the mix of free amino acids. The total free amino acid mixes for the UA and UB diets were adjusted to 7.1% of the total dry diet (Table 1) by adding more non-essential amino acids.

|  |  | Diet |  |  |  |
| --- | --- | --- | --- | --- | --- |
|  | Amino acid (g/kg) | REF | B | UA | UB |
| Essential | Arginine | 12.87 | 9.64 | 9.64 | 9.64 |
|  | Histidine | 5.02 | 3.76 | 3.76 | 3.76 |
|  | Isoleucine | 9.05 | 6.78 | 6.78 | **3.61** |
|  | Leucine | 15.56 | 11.65 | **6.21** | 11.65 |
|  | Lysine | 11.06 | 8.28 | 8.28 | **4.41** |
|  | Methionine | 4.41 | 3.3 | 3.3 | 3.3 |
|  | Phenylalanine | 9.79 | 7.33 | **3.91** | 7.33 |
|  | Threonine | 7.49 | 5.61 | **2.99** | 5.61 |
|  | Tryptophan | 2.32 | 1.74 | 1.74 | **0.93** |
|  | Valine | 10.18 | 7.62 | 7.62 | 7.62 |
| Non-essential | Alanine | 9.22 | 6.9 | 7.82 | 7.53 |
|  | Asparagine | 5.30 | 3.97 | 4.5 | 4.33 |
|  | Aspartic acid | 18.90 | 14.15 | 16.03 | 15.44 |
|  | Cysteine | 3.23 | 2.42 | 2.74 | 2.64 |
|  | Glutamine | 5.30 | 3.97 | 4.5 | 4.33 |
|  | Glutamic acid | 36.53 | 27.35 | 30.99 | 29.84 |
|  | Glycine | 8.53 | 6.39 | 7.23 | 6.97 |
|  | Proline | 11.53 | 8.63 | 9.78 | 9.41 |
|  | Serine | 10.26 | 7.68 | 8.7 | 8.38 |
|  | Tyrosine | 6.57 | 4.92 | 5.57 | 5.37 |

### Growth analysis

To follow the daily growth, 15 larvae were each day randomly selected and weighed to determine the average mass per replicate, after which they were returned to the feed. At the end of the experiment, surviving larvae were counted and weighed, and mortality was calculated. Daily total larval mass was estimated by multiplying the mean larval mass by the estimated number of live larvae each day, under the assumption that mortality was gradual over the seven-day period. These estimates facilitated the tracking of daily growth and its relation to daily gas exchange and metabolic rate. The growth progression is also evaluated using a specific growth rate (SGR), calculated as follows:

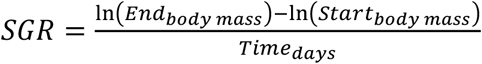

### Water, lipid, and protein content

Following the experimental period, all larvae from each replicate were oven-dried at 60°C for 3-4 days until a stable mass was reached. Water content was determined by subtracting dry mass from wet mass, and from each replicate, 40 larvae were subsampled to assess lipid and protein contents. For lipid extraction, dried larvae were weighed and then washed five times with petroleum ether, where each wash was followed by a 48-hour soak to ensure thorough lipid dissolution. The defatted samples were then re-dried in the oven at 60°C until they had a stable mass, and crude lipid content was calculated as the difference between initial and lean dry masses. Finely ground defatted sub-samples (4-7 mg) underwent nitrogen analysis using a combustion analyser (Na 2000, Italy). Crude protein content was then estimated as the product of the nitrogen content (%) and 4.76 g protein/g nitrogen, which is an established conversion factor for the protein-to-nitrogen ratio in BSFL (Janssen *et al*., 2017).

### Gas exchange measurements

An open-flow respirometry setup generally followed the protocol described in Schøn *et al*. (2025a, Schøn *et al*. 2025b) with custom-built, airtight respiration chambers (15 cm diameter, 6 cm height, 1 L volume) placed in a temperature-controlled cabinet set at 27 °C. A continuous flow of room air (~600 mL/min) passed through each chamber, ensuring a stable O_2_ supply while removing water vapour and CO_2_. The input air flow to each chamber was continuously measured using a Flowbar-8 Mass Flow Meter, Sable Systems, and the dry input flow of air (V_I_) was recalculated to flow of standard dry air (STPD) assuming room air had a temperature of 21 °C, 50% RH and atmospheric pressure of 760 mmHg.

To measure gas exchange, a sub-sampler pump (SS-4 Sub-Sampler Pump, Sable Systems) and an 8-channel multiplexer (RM-8 Respirometry Flow Multiplexer, Sable Systems) sequentially sampled air (~200 mL/min) from each of the eight chambers (seven experimental and one empty control). The sampled air passed through a calcium chloride column to remove water vapour and a syringe filter to remove any particles (Acrodisc syringe filter, 25 mm, 0.2 µm Supor membrane, Pall Corporation) before entering the gas analysers. The O_2_ and CO_2_ concentrations were measured using a differential O_2_ analyser (FC-2 Differential Oxygen Analyser (Oxzilla), Sable Systems) and a CO_2_ analyser (LI-850 Gas Analyzer, LI-COR Biosciences) that were routinely calibrated against known gas mixtures. The multiplexer cycled between chambers every 7.5 minutes, enabling hourly measurements of CO_2_ production and O_2_ consumption from each chamber throughout the 7-day experiment. Recordings from each chamber of flow, CO_2_ fraction and O_2_ fraction were allowed to stabilise during the 7.5 min period and only the last 30 sec of data was used to assess fractional gas content and flow (see Schøn *et al*. (2025b) for example traces in a similar set up). The hourly measurement of fractional CO_2_ and O_2_ content measured in the empty chamber was used for the assessment of gas fractions in input air (F_I_O_2_ and F_I_CO_2_, respectively) while output fractions were measured from each experimental chamber hourly (F_E_O_2_ and F_E_CO_2_, respectively).

The rate of oxygen consumption 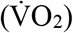 and carbon dioxide production 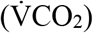 (mL/min) were found by multiplying calculated input of dry air (V_I_) (mL/min) with the difference in gas concentrations between experimental (output) and control chambers (input). Because input and output volumes are not identical when RER is different from 1, we corrected for the difference in outflow volume as outlined in Withers (2001). Specifically, 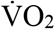 was calculated as:

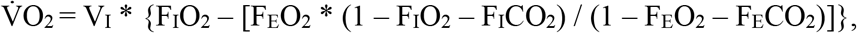

and 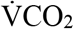 was calculated as:

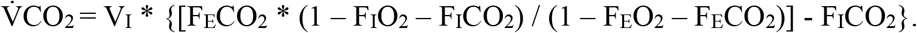

For the hourly estimates of 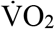 and 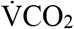 we would sometime experience unstable measurements of either flow or fractional gas content. Such outliers were removed, and the missing measurements were replaced with values interpolated from prior and subsequent high-quality measurements using the data quality assessment protocol described by Schøn *et al*. (2025b). Total gas exchange (VO_2_ and VCO_2_) (mL) was calculated for each replicate across the 7-day growth period by summation of all hourly measurements. These values were used to determine the temporal 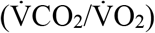 and cumulative (VCO_2_/VO_2_) respiratory exchange ratio (RER). Over the 7-day experiment, each chamber lost an average of 28.7 ± 2.5 g per day, mainly due to evaporation and CO_2_ production.

It should be noted that we could unfortunately not control for the metabolic contribution from microbial activity. Although microbial respiration likely occurs in our system, we content that the overall metabolic contribution from fungi and bacteria is likely limited compared to that of the growing larvae when evaluated after seven days. It is unfortunately not possible to use chambers with feed and no larvae as control as our pilot experiments revealed gas exchange patterns from such chambers that were completely different from conditions where larvae are present as larval presence hugely affects microbial respiration (see also discussion).

### Data Analysis

One-way ANOVA was used to assess differences in BSFL growth, gas exchange, and body composition across treatment groups, following confirmation of normality and variance homogeneity. Tukey’s Honest Significant Difference (HSD) test was applied for post hoc comparisons, with significance set at α = 0.05. A linear regression was applied across dietary treatments to describe the relationships between biomass and O_2_ production and between energetic gains and expenditures. Data processing and visualisation were conducted in RStudio (version 2024.04.2, R version 4.3.2) (Posit Team, 2023; R Core Team, 2023).

## Results

### Growth and body composition

All experimental diets supported larval growth and development with mean survival rates ranging from 88–99%, although larvae on the UB diet exhibited a marginal but statistically significant increase in mortality (Table 3). The REF diet resulted in the highest total dry matter yield per replicate (32.4 g from ~ 600 larvae; Fig. 1A; Table 3). Total larvae yield in the B diet (24.7 g) was lower than that of larvae reared on the REF diet, but higher than for larvae on the two unbalanced diets, UA (16.9 g) and UB (12.1 g) (Tukey HSD, *α* = 0.05; insert in Fig. 1A). In accordance with the higher total growth, the average SGR was higher on the REF and B diets than in UA and UB (Tukey HSD, α = 0.05; insert in Fig. 1B, Table 3). It should be noted that larvae on the REF diet did not grow after reaching day five to six. This indicates that larvae on the REF diet had reached the prepupal wandering stage (Gligorescu *et al*. 2019; Schøn *et al*. 2025b), whereas larvae on the other experimental diets were still generally gaining mass on day seven when measurements ended (Fig. 1) indicating that they were still in the growth phase.

**Table 3.** Survival, growth, body composition, and gas exchange of black soldier fly larvae reared on diets with variable amino acid profiles (Table 2).

|  | Chicken feed<br>(REF) | Balanced<br>(B) | Unbalanced A<br>(UA) | Unbalanced B<br>(UB) |
| --- | --- | --- | --- | --- |
| **Growth performance** |  |  |  |  |
| Survival rate (%) | 99.0 ± 1.5%<sup>a</sup> | 93.9 ± 4.0%<sup>a</sup> | 92.7 ± 3.9%<sup>a</sup> | 88.1 ± 7.6%<sup>b</sup> |
| Larvae dry mass (DM, g per replicate) | 32.4 ± 1.0 g<sup>a</sup> | 24.7 ± 3.9 g<sup>b</sup> | 16.9 ± 3.1 g<sup>c</sup> | 12.1 ± 4.0 g<sup>c</sup> |
| Specific growth rate (per day) | 0.50 ± 0.06<sup>a</sup> | 0.47 ± 0.05<sup>a</sup> | 0.39 ± 0.04<sup>b</sup> | 0.36 ± 0.04<sup>b</sup> |
| Larvae dry mass (% DM) | 30.8 ± 1.1%<sup>a</sup> | 31.2 ± 1.4%<sup>a</sup> | 33.5 ± 1.6%<sup>b</sup> | 33.3 ± 1.1%<sup>b</sup> |
| Larvae crude protein (% of DM) | 34.9 ± 1.5%<sup>a</sup> | 29.0 ± 1.7%<sup>b</sup> | 24.7 ± 1.3%<sup>c</sup> | 23.4 ± 1.4%<sup>c</sup> |
| Larvae crude lipid (% of DM) | 10.7 ± 2.5 %<sup>a</sup> | 26.1 ± 2.9 %<sup>b</sup> | 40.5 ± 3.1%<sup>c</sup> | 37.4 ± 3.4%<sup>c</sup> |
| **Gas exchange measurements** |  |  |  |  |
| VCO<sub>2</sub> (L, total for 7 days) | 34.8 ± 0.77 L<sup>a</sup> | 21.5 ± 1.29 L<sup>b</sup> | 16.5 ± 1.17 L<sup>c</sup> | 11.3 ± 1.56 L<sup>d</sup> |
| VO<sub>2</sub> (L, total for 7 days) | 32.2 ± 1.25 L<sup>a</sup> | 15.3 ± 1.08 L<sup>b</sup> | 10.4 ± 0.71 L<sup>c</sup> | 7.39 ± 0.97 L<sup>c</sup> |
| RER | 1.09 ± 0.032<sup>a</sup> | 1.42 ± 0.028<sup>b</sup> | 1.58 ± 0.054<sup>c</sup> | 1.53 ± 0.38<sup>bc</sup> |

RER = respiratory exchange ratio. RER is calculated from total VCO<sub>2</sub>/VO<sub>2</sub>. Values are mean ± standard error. Different letters indicate significant differences (one-way ANOVA followed by Tukey HSD test, $\alpha = 0.05$ , n = 7). DM = dry matter, V = volume.

**Figure 1.**
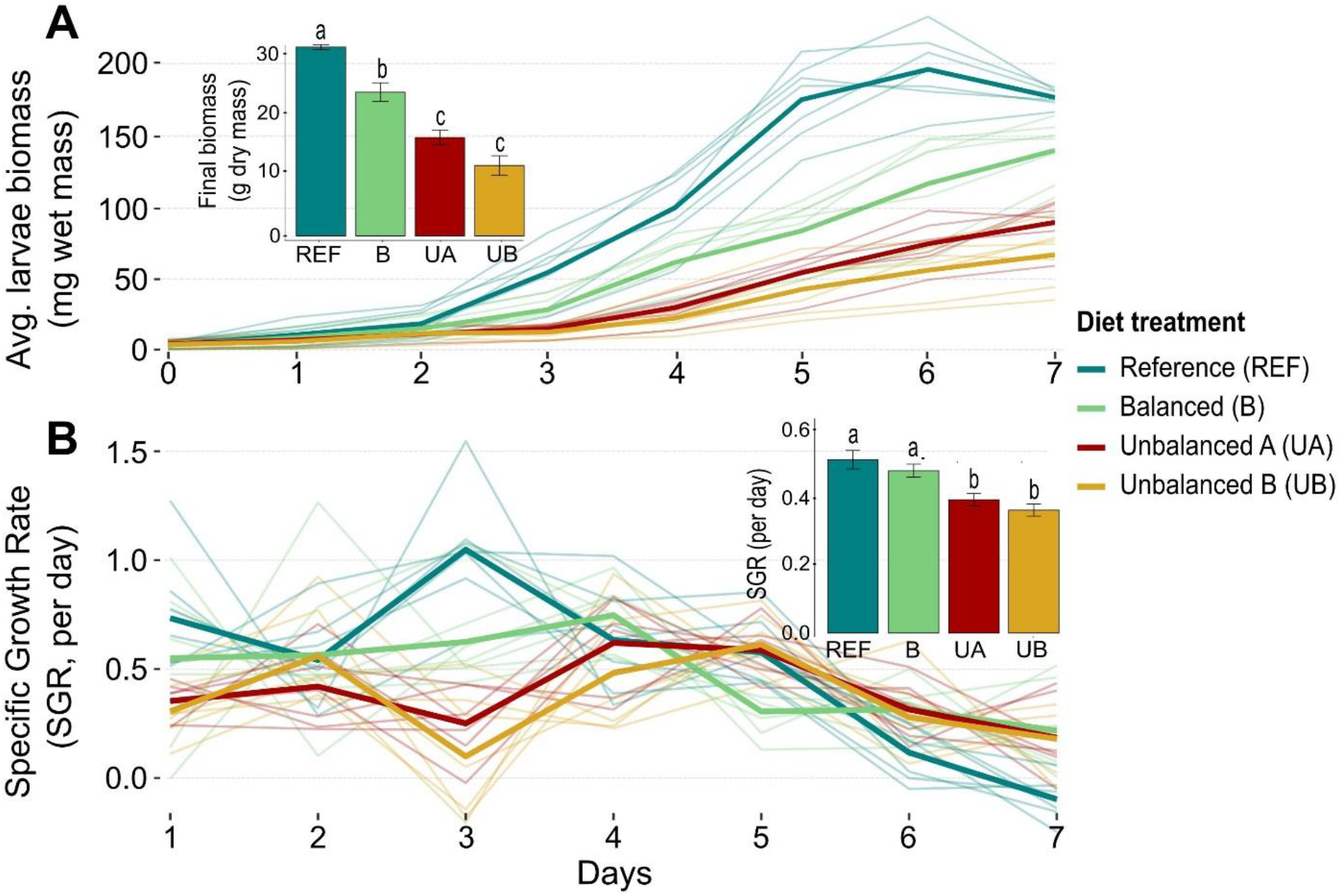
Dietary effects on growth of black soldier fly larvae during a 7-day rearing period. A) Temporal changes in average individual larval mass (mg wet mass), with insert depicting final total larval biomass (dry mass) per treatment. B) Daily estimates of specific growth rate (SGR), based on the change in estimated larval wet mass in a replicate between consecutive days. The B) insert shows an average SGR over the full 7-day period for each treatment. Bold lines represent treatment means and semi-transparent lines indicate individual replicates (n = 7). Insert bar graphs show means ± SE, where distinct letters denote statistically significant differences among treatments (one-way ANOVA with Tukey’s HSD, α = 0.05).

Analysis of larval body composition revealed pronounced diet-dependent differences in lipid and protein content of the larvae after the 7-day rearing period. Crude protein content in the dry mass was highest in larvae reared on the REF diet (34.9%), intermediate in larvae reared on the B diet (29.0%), and lowest in larvae reared on the two unbalanced diets, UA(24.7%) and UB (23.4%) (Table 3). Conversely, crude lipid content was lowest in larvae from the REF diet (10.7%), more than twice as high in larvae from the B diet (26.1%), and almost four times as high in larvae from the unbalanced diets, UB (37.4%) and UA (40.5%) (Table 3).

### Gas exchange rates (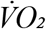 and 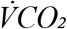)

Oxygen consumption rate 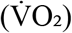 and carbon dioxide production rate 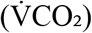 were measured hourly over the 7Cday experimental period. All treatment groups were initially characterised by parallel increases in 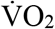 and 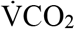 (Fig. 2) with distinct peaks over the experiment, but the absolute rates and temporal developments of gas exchange differed markedly among treatments (Fig. 2). Larvae reared on the REF diet exhibited 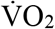 peaks around day 3.5 (4.5 mL O_2_/min) and 5.5 (10 mL O_2_/min) followed by a marked reduction in gas exchange rate on day 6 and 7, matching the time when these larvae had stopped growing (Fig. 1). Larvae on the B diet followed a similar pattern, with slightly delayed peaks (day 4 and day 6.5) and more modest absolute gas exchange rates. The two semi-artificial, unbalanced diets showed even lower gas exchange rates with further deductions in temporal development and peak amplitudes. Thus, in the two unbalanced treatments, gas exchange peaked around day 5 (1.3 and 1.9 mL O_2_/min, respectively) with a second rise emerging near the end of the experiment on day 7. When integrating the oxygen consumption over the entire period (insert in Fig. 2A), the total VO_2_ was higher on the REF diet (32.2 L O_2_) compared to the B diet (15.3 L O_2_). Likewise, the total VO_2_ in the B diet was higher than both in UA (10.4 L O_2_) and UB (7.4 L O_2_) (Tukey HSD, α = 0.05).

**Figure 2.**
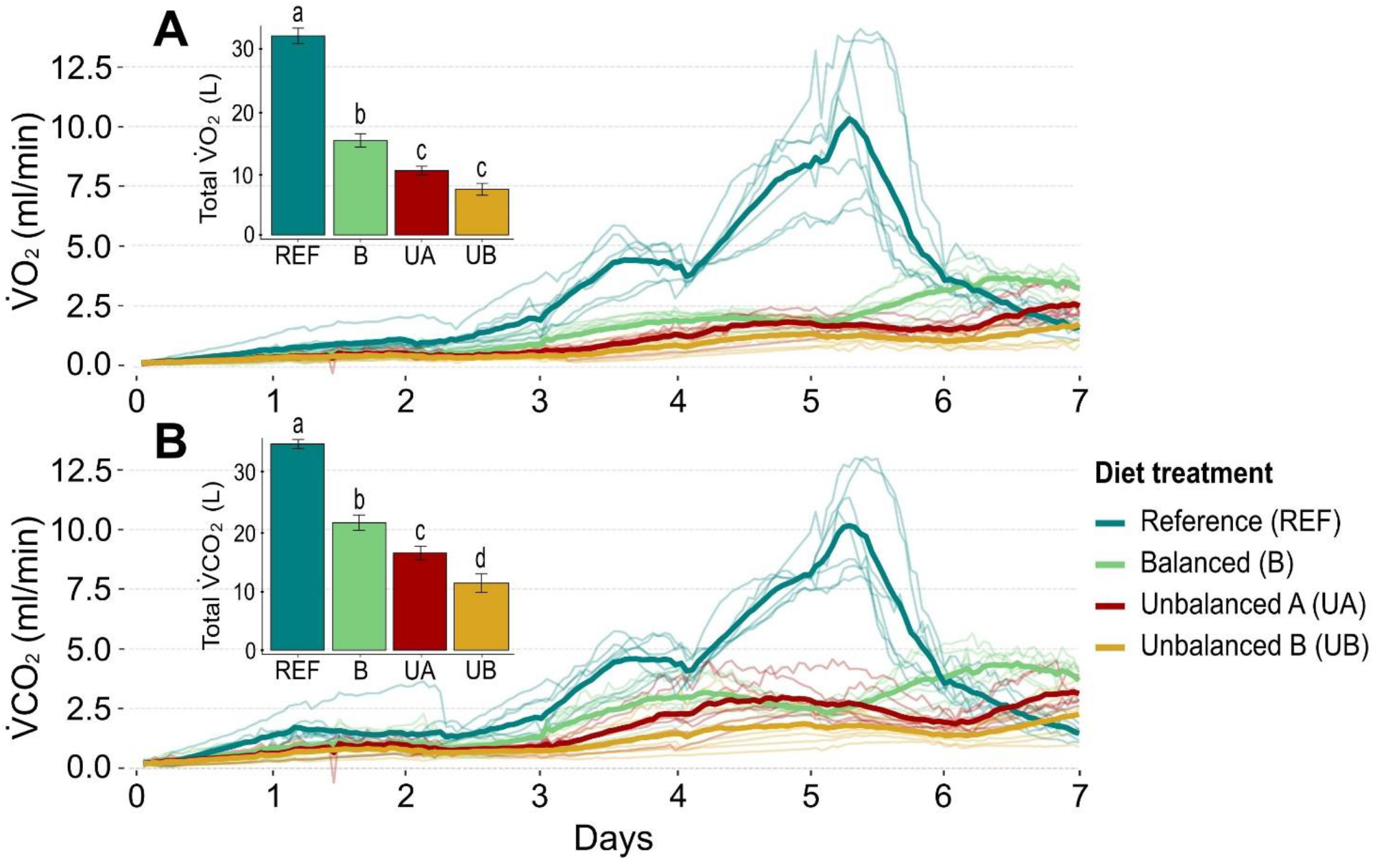
Dietary effects on gas exchange in black soldier fly larvae (BSFL) during a 7-day rearing period. Temporal trajectory of A) oxygen consumption rates 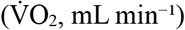 and B) carbon dioxide production rates 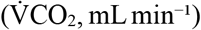, with insert summarising total gas exchange (VO_2_ and VCO_2_, L) across the full experimental duration for each diet treatment. Bold lines represent treatment means and semi-transparent lines indicate individual replicates (n = 7). The insert bar graphs show means ± SE, where distinct letters denote statistically significant differences among treatments (one-way ANOVA with Tukey’s HSD, α = 0.05).

In the present study, all diet treatments were initially characterised by a RER around 1.5-2.0, which declined gradually over the 7-day period. The gradual decrease in RER was faster and more pronounced in the REF diet (Fig. 3), and there was a tendency for the semi-artificial balanced diet (B) to have slightly lower RER than the two semi-artificial unbalanced diets (UA and UB). When cumulative gas exchange was integrated over the entire 7-day growth period, we found that larvae on the REF diet had a lower RER (1.09). The B diet had a cumulative RER of 1.42 which was slightly lower, but not significantly different from the RER values of the two unbalanced diets (UA: 1.58; UB: 1.53) (Tukey HSD, α = 0.05; Fig. 3 insert).

**Figure 3.**
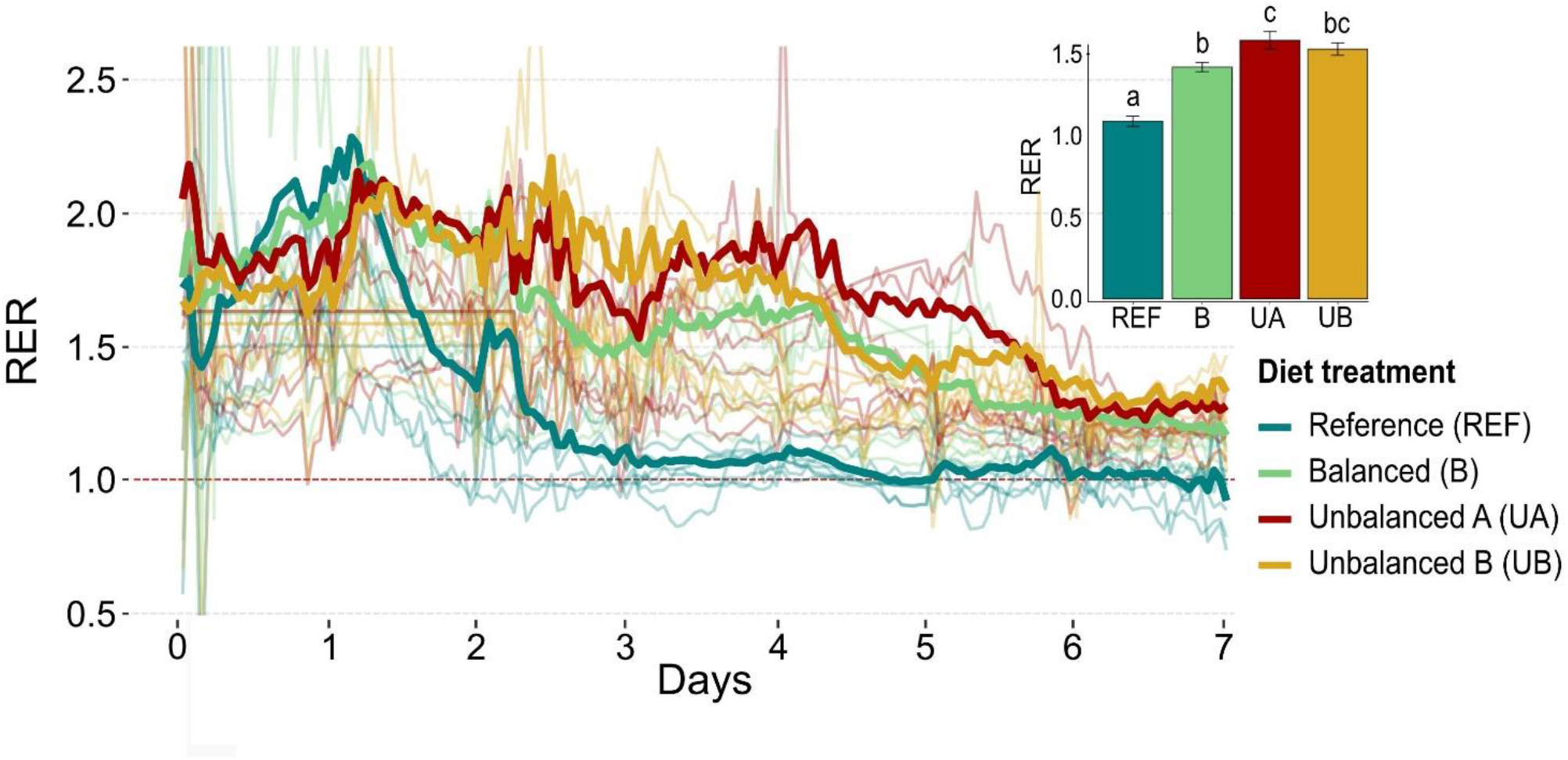
Respiratory exchange ratio (RER; 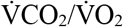) for black soldier fly larvae (BSFL) during a 7-day rearing period plotted over time. Insert bar showing RER derived from cumulative O_2_ consumption and CO_2_ production per treatment (VCO_2_/VO_2_). Bold lines represent treatment means and semi-transparent lines indicate individual replicates (n = 7). The insert bar graph shows means ± SE, where distinct letters denote statistically significant differences among treatments (one-way ANOVA with Tukey’s HSD, α = 0.05).

### Energetic cost of growth and protein assimilation

Integrating growth, body composition, and metabolic rate allows evaluation of the total energetic cost of growth by relating oxygen consumption to energy stored in biomass, with protein and lipid deposition expressed as energy equivalents. When analysing all treatment diets, we found a strong correlation (R^2^ = 0.83) between the total larval dry mass and total O_2_ consumption across individual replicates. Thus, on average each g of dry biomass assimilation was associated with the use of 1.08 L O_2_ for aerobic metabolism (Fig. 4A). When this analysis was performed for each treatment, we found the apparent “gross cost of growth” to be higher for larvae on the REF diet (1.00 L/g), while larvae on the three treatment diets all had lower cost of growth around 0.62 L/g (Tukey HSD, α = 0.05; insert in Fig. 4A).

**Figure 4.**
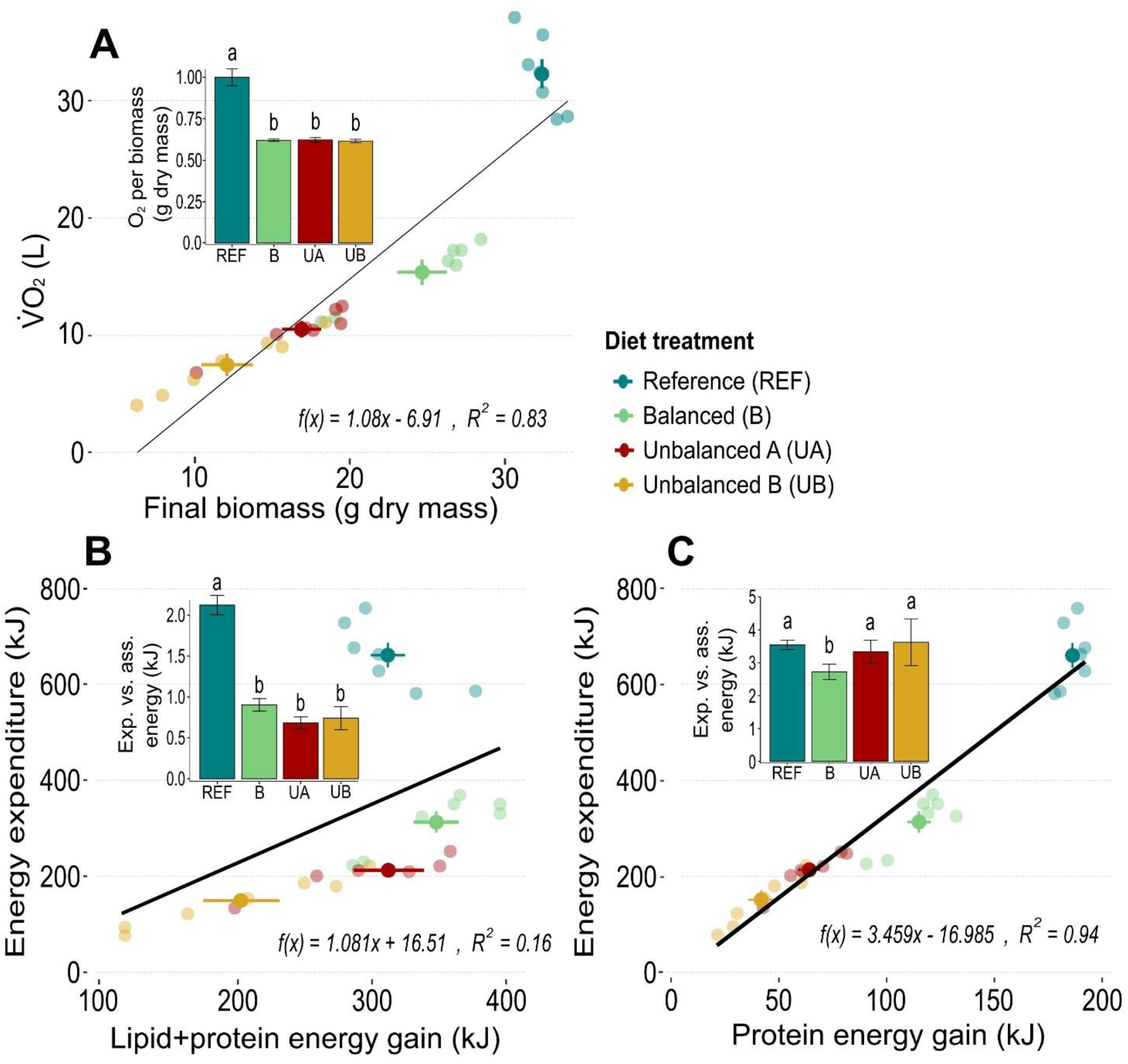
Metabolic costs of growth during a 7-day rearing period in black soldier fly larvae. A) Linear relationship between total oxygen consumption (L) and final larval dry biomass (g). B) and C) Energy expenditure of the system, calculated by multiplying VO_2_ by the caloric equivalent of oxygen (20.5 kJ L^−1^), plotted against assimilated energy in the biomass (kJ), represented as gain in lipid plus protein (B) or protein alone (C). Bold points represent treatment means and semi-transparent points show individual replicates (n = 7). Black lines representing linear regression models for all replicates (n = 28). The insert bar graphs show means ± standard error for the ratio between the expended energy (Exp.) and the assimilated energy (Ass.), where distinct letters denote statistically significant differences among treatments (one-way ANOVA with Tukey’s HSD, α = 0.05).

To evaluate a more direct ratio of metabolic energy expenditure to assimilated energy, we converted the cumulative O_2_ consumption to metabolic energy using a standard oxy-calorific coefficient of 20.5 kJ L O_2_^−1^ (Schmidt-Nilsen, 1997). Likewise, we calculated assimilated energy from the increment of lipid and protein by multiplying the measured fractions by their respective energy densities (37 kJ g^−1^ for lipid and 17 kJ g^−1^ for protein) (Schmidt-Nilsen, 1997). As seen in Fig. 4B, this analysis revealed marked differences in the energetic cost of energy assimilation among treatments and accordingly the general association between energy expenditure and energy assimilation had a weak correlation among all replicates (R^2^ = 0.16). When analysed within replicates, we found that the costs of energy assimilation were more than twice as high in the REF treatment compared to the three semi-artificial diets (Fig. 4B, insert). Accordingly, larvae fed on the REF diet expended 2.12 kJ per kJ lipid and protein assimilated. In contrast, larvae fed on the semi-artificial diets showed no significant differences, with the balanced diet yielding an expenditure of 0.90 kJ, and the unbalanced diets 0.68 kJ (UA) and 0.73 kJ (UB) per kJ lipid and protein assimilated (Fig. 4B, insert) (Tukey HSD, α = 0.05).

Restricting the analysis to consider only the cost of protein assimilation we found a strong and general correlation (Fig 4C; R^2^ = 0.94) with a slope of 3.46 kJ energy expended per kJ protein assimilated. Despite the strong general correlation, the energetic cost of protein assimilation also differed among diets (Fig. 4C, insert). Larvae fed the semi-artificial balanced diet (B) showed the lowest cost (2.73 kJ), while REF (3.55 kJ) and UA (3.34 kJ) showed intermediate costs, and UB was the least efficient, with a cost of 3.63 kJ per kJ protein assimilated) (Tukey HSD, α = 0.05).

## Discussion

By combining respirometry with chemical analysis, we quantified how suboptimal dietary EAA profiles affects growth and metabolic expense in BSFL. The primary aim was to compare a semi-synthetic diet with balanced EAA composition with two other semi-synthetic diets with the same macronutrient composition, but with two different unbalanced EAA profiles.

### Growth, body composition, and survival

Survival was slightly but significantly reduced on the UB diet compared to the REF diet (Table 3), indicating that nutritional imbalance can potentially compromise viability. The BSFL generally tolerate wide ranges of dietary protein (Berggreen *et al*., 2025; Oonincx *et al*., 2015; Schneider *et al*., 2025; Schøn *et al*., 2025a), and high survival is also reported under isonitrogenous diets with imbalanced EAA profiles (Berggreen *et al*., 2025). Nonetheless, severe EAA deficiencies or excesses can elevate mortality in BSFL (Koethe *et al*., 2022; Spranghers *et al*., 2025), as also seen in medflies (Davis, 1975).

We found that unbalanced EAA diets markedly reduced growth of BSFL compared to the balanced diet (Table 3, Fig. 1). Biomass accumulation is known to depend on the total dietary protein content, and optimal macronutrient ratios have been widely studied (Barragan-Fonseca *et al*., 2019; Berggreen *et al*., 2025; Broeckx *et al*., 2021, Schøn *et al*. 2025a, Eggink *et al*. 2023). However, micronutrient requirements, particularly requirements for individual EAAs, remain poorly defined for BSFL (Tomberlin *et al*., 2023). In the present study, combined restriction of three EAAs resulted in impaired growth, in accordance with recent findings showing that growth was also suppressed when individual EAAs were deficient (Berggreen *et al*., 2025). In contrast, Lemme and Klüber (2024) reported no effects on growth following individual EAA reductions. Such discrepancies may reflect masked treatment effects due to uncontrolled EAA or micronutrient levels in the basal diets.

Larvae reared on unbalanced diets were characterised by lower fractional protein content and higher fractional lipid content than those fed on the B or REF diet (Table 3). This is consistent with effects of diets deficient in only a single EAA (Berggreen *et al*., 2025) and is likely reflecting a metabolic prioritisation of the growing larvae. Specifically, protein synthesis is prioritised over fat storage (Lee, 2007), especially in early life stages (Talal *et al*., 2024), when amino acids are adequate and balanced (Lee, 2007). However, when the amino acid composition is unbalanced, excess non-limiting amino acids are likely deaminated and some of the carbon is stored as lipids (Stryer, 1995). Such changes in metabolism could explain why larvae on unbalanced diets show the highest fractional lipid content. Lipid accumulation in larvae on unbalanced diets may also result from compensatory feeding under nutrient limitation (Raubenheimer and Simpson, 1997) leading to excess intake of protein and carbohydrate subsequently converted to lipid. Similar mechanisms are reported in other insects (Berner *et al*., 2005; Lee, 2007) and have been proposed for BSFL (Banks *et al*., 2014). This is further supported by Pimentel *et al*. (2017) who reported that, in BSFL, protein-poor diets downregulate genes involved in protein storage and promote lipid accumulation via de novo lipogenesis.

### Gas exchange

Gas exchange patterns revealed distinct metabolic responses to dietary amino acid balance (Fig. 2). Both oxygen consumption (VO_2_) and carbon dioxide production (VCO_2_) were reduced in larvae fed the UA and UB diets (Fig 2 inserts) with suppressed respiratory peaks and delayed development relative to the B diet (Fig 2). Similar patterns were reported by Schøn *et al*. (2025a) under low protein-to-carbohydrate conditions, which may also be considered low-quality diets. The reduced gas exchange likely reflects the difference in biomass as more larval biomass results in higher metabolic turnover for maintenance metabolism, but differences are likely also related to lower energetic demand due to suppressed anabolic activity associated with protein synthesis and tissue growth. In BSFL, respiratory metabolism scales with growth performance (Parodi *et al*., 2020), and the observed metabolic downregulation is consistent with patterns seen under other growth-limiting conditions, such as low dietary water content in *Manduca sexta* (Van’t Hof and Martin, 1989) or thermal stress in BSFL (Schøn *et al*., 2025b). The developmental delay may reflect prolonged larval instars, where moulting phases following respiratory peaks are associated with reduced feeding (Gligorescu *et al*., 2019). The substantial elevated gas exchange in REF fed larvae, also reported by Schøn *et al*. (2025a), likely reflects greater availability of growth-promoting micronutrients in the REF diet that enhance anabolic processes.

When calculated from the total CO_2_ and O_2_ exchange over the 7-day period, the RER exceeded 1.00 across treatments. The highest RER values were observed in larvae fed the B, UA, and UB diets, with an apparent positive association with crude lipid content across treatments (Table 3). While CO_2_ was not used to estimate cost of growth (see below), it was used to calculate RER, to get information about the combined impact of metabolic respiratory activity and the surplus CO_2_ produced during lipogenesis (Talal *et al*., 2021). Accordingly, elevated RER values suggest that CO_2_ production arose not only from catabolism (RER: 0.7–1.0, depending on lipid or carbohydrate utilisation, respectively (Schmidt-Nilsen, 1997)), but also from anabolic lipogenesis (Schøn *et al*., 2025b; Talal *et al*., 2021) and possibly microbial metabolism during early development (Parodi et al., 2020). This supports the suggestion by Berggreen *et al*. (2025) that impaired protein accretion redirects excess amino acids toward lipid synthesis (Berggreen *et al*., 2025), increasing lipid synthesis and elevating CO_2_ production relative to O_2_ consumption. However, this finding was expected, as RER values above 1 are common among fast-growing animals (Jakobsen and Thorbekt, 1993; Talal *et al*., 2021) including BSFL (Parodi *et al*., 2020; Schøn *et al*., 2025a, 2025b). As we measured gas exchange of larvae including the putative microbial metabolism from the feed, it is not possible to separate larval from microbial metabolism, and some of the excess CO_2_ measured likely originates from microbial fermentation (Fuhrmann *et al*. 2025; Parodi *et al*. 2020). In pilot experiments we investigated microbial gas exchange in the feed after removing larvae, and while this gas exchange was initially limited it increased dramatically within hours. This shows that in the absence of larvae the metabolism from microorganisms quickly becomes much higher than when larvae are present, probably because larvae keep microorganism numbers low, and the contributions from larvae and microorganisms can therefore not be separated by including treatments without larvae. We here treat the metabolism of the system as a whole with the contention that the main driver of respiration originates from the larvae and their growth.

### Energetic cost of growth and protein assimilation

The analysis of gross cost of growth was evaluated by relating total oxygen consumption to biomass gain. Cumulative oxygen consumption over the 7-day period was converted to metabolic energy using a standard oxycaloric coefficient, assuming metabolism primarily fuelled by mixed nutrients. Biomass gain was expressed as the energy stored in newly deposited protein and lipid, calculated from their respective energy densities. This method integrates metabolic costs as cumulative output relative to biochemical gain and avoids potential biases associated with CO_2_-based estimates, which, as described above, can be confounded by lipogenesis and microbial activity (Gold *et al*., 2018; Schøn *et al*., 2025b; Talal *et al*., 2021). The gross cost of growth reflects the combined demands of tissue synthesis, maintenance metabolism, and microbial processes (Eriksen, 2022; Laganaro *et al*., 2021; Fuhrmann *et al*., 2025), and while our whole-system measurements do not separate these components, relative differences in cost per unit growth still allow meaningful comparisons of substrate efficiency and nutrient limitations.

As an indirect measurement of energy metabolism, the total oxygen consumption scaled linearly with final larval biomass (Fig. 4A), reflecting the well-established link between total feed assimilation (growth) and energy consumption in insects (Laganaro *et al*., 2021; Von Bertalanffy, 1957). The gross cost of growth (volume oxygen used per gram dry body mass gained) remained consistent across all three treatment diets (Fig. 4A, insert) demonstrating that the metabolic energy used to produce a g of biomass on these three diets did not differ. Larvae fed on the REF diet achieved the greatest biomass accumulation but also exhibited higher gross cost of growth per gram of biomass assimilated than those on semi-synthetic diets. Notably, larvae on the REF diet showed a distinct temporal growth pattern, characterised by ~1.5 days of biomass loss and reduced metabolic rate near the end of the experiment, presumably because the average larvae had reached the non-feeding prepupal larval stage. Oxygen consumed during this period was primarily allocated to catabolic or maintenance processes rather than net growth, potentially overestimating the true cost of tissue synthesis. A more precise definition of larval stages would allow for a more detailed analysis of growth costs. Nonetheless, larvae on the three semi-artificial diets remained in the growth phase throughout the 7-day experiment and are therefore evaluated on more equal terms.

To understand why mass-specific metabolic costs remained unchanged between the balanced (B) and unbalanced diets (UA and UB), we assessed energy expenditure relative to total energy gain (lipid + protein energy) (Fig. 4B). Again, energetic costs appeared consistent across diets, likely due to differences in biomass composition. Protein synthesis is more metabolically demanding than lipid accumulation (Hawkins, 1991; Secor, 2009) and likely constitutes a larger component of energy expenditure in larvae fed the B and REF diets, which best supported protein growth. In contrast, larvae on EAA-deficient diets face suboptimal conditions for growth due to limited substrates for protein synthesis. As a result, assimilated nutrients are redirected toward lipid storage, and the relative allocation of energy shifts so that growth is constrained while maintenance becomes relatively more dominant (Laganaro *et al*., 2021). Although total protein synthesis is reduced in larvae fed unbalanced diets, the metabolic costs of lipid storage, maintenance, and deamination of excess amino acids (Hawkins, 1991) may raise energy expenditure to levels comparable to protein-synthesising larvae.

The relatively high energetic cost of protein assimilation is supported by comparison of the weak association between total energy gain (lipid + protein) and energy expenditure (Fig. 4B), in contrast to the stronger correlation observed for protein gain versus energy expenditure (Fig. 4C). These results suggest that, while lipid accumulation incurs some energetic cost, protein synthesis is the primary driver of energy expenditure during growth. This aligns with the prediction of Eriksen (2022), who modelled BSFL energy budgets using CO_2_-based fluxes and partitioned respiration into maintenance and biosynthesis. Despite methodological differences, both approaches converge on the conclusion that protein synthesis is substantially more energetically costly than lipid deposition and represents the dominant metabolic driver in BSFL. Thus, when focusing solely on protein gain, the energetic inefficiency of EAA-deficient diets became evident. Larvae on UA and UB expended more energy per kJ of protein deposited than those on the B diet (Fig. 4C), and the same was observed for larvae fed the REF diet, likely due to the inclusion of energy expenditure during the post-feeding prepupal stage in the total energy calculation. As diets with unbalanced EAA composition caused lower protein deposition and higher energetic cost of protein growth, our results indicate that suboptimal dietary profiles may constrain the efficiency of insect production systems, highlighting the need for further research to improve the sustainability and economic viability of insect feeds.

## Conclusions

This study demonstrates that unbalanced EAA profiles impair growth performance and metabolic efficiency in BSFL. By integrating growth, body composition, and gas exchange data, we show that suboptimal EAA profiles limit protein synthesis, promote lipid accumulation, and increase the gross energetic cost of amino acid metabolism and protein deposition. Metabolic costs per unit of protein deposited were substantially higher on EAA-deficient diets (up to 3.63; kJ per kJ protein) than on the balanced diet (2.73 kJ per kJ protein), reflecting inefficient amino acid utilization. Whether unbalanced diets are disadvantageous from a feed-cost perspective depends on the trade-off between reduced larval growth and protein assimilation versus the potentially lower cost of low-quality feed ingredients. The use of crystalline amino acids offers precise control of dietary composition, but their viability depends on whether they support sufficient biomass yield and quality. Our results suggest that protein from complex feed sources (e.g. chicken feed), may be more efficiently assimilated than crystalline amino acids, even when EAA profiles are matched, which is also previously reported (Berggreen *et al*., 2025). We here included deficient amounts of three EAAs per unbalanced diet to ensure that we would find metabolic effects. Theoretically, the single most limiting EAA alone caused the measured effect within each unbalanced diet, which could be tested in future studies. By quantifying the physiological costs of EAA imbalance, this study provides benchmarks to support the evaluation of suboptimal feeds in BSFL production. Further research should assess whether targeted EAA supplementation can improve protein yield and energy efficiency on low-quality substrates, thereby contributing to the development of nutritionally and economically optimized insect feeds.

## Acknowledgements

This study was supported by the Danish Green Development and Demonstration Programme (GUDP): EntoFeed 34009-21-1837 and FlyCloud 34009-20-1652.

## Supplementary Information

**Table S1.**
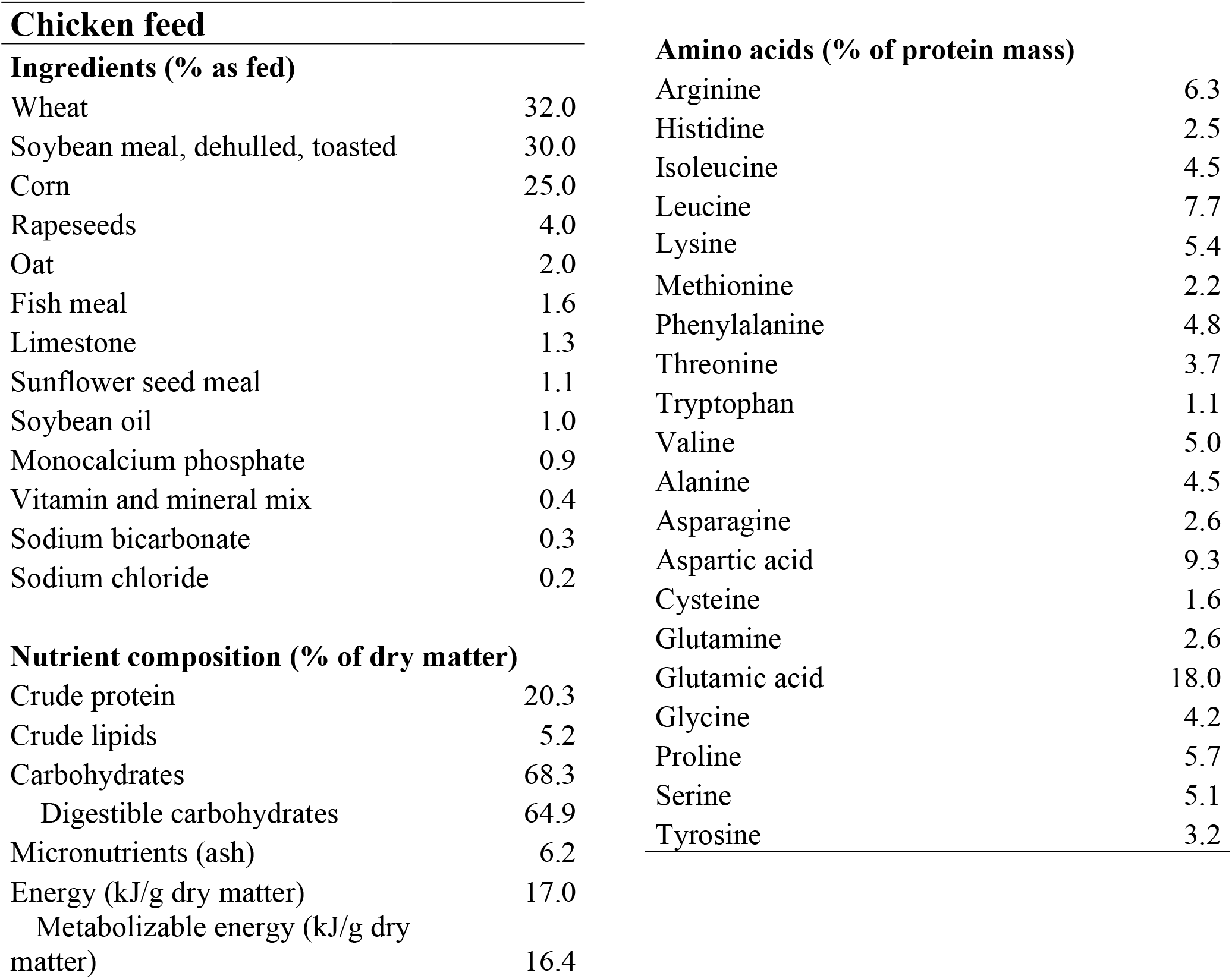
Ingredient and nutrient compositions in the chicken feed (Kyllekræs 1, Danish Agro) used in the experiment. Ingredient and nutrient composition were given by the producer, and amino acid contents were analysed.

**Ingredients (% as fed)**
|  |  |
| --- | --- |
| Wheat | 32.0 |
| Soybean meal, dehulled, toasted | 30.0 |
| Corn | 25.0 |
| Rapeseeds | 4.0 |
| Oat | 2.0 |
| Fish meal | 1.6 |
| Limestone | 1.3 |
| Sunflower seed meal | 1.1 |
| Soybean oil | 1.0 |
| Monocalcium phosphate | 0.9 |
| Vitamin and mineral mix | 0.4 |
| Sodium bicarbonate | 0.3 |
| Sodium chloride | 0.2 |
| **Nutrient composition (% of dry matter)** |  |
| Crude protein | 20.3 |
| Crude lipids | 5.2 |
| Carbohydrates | 68.3 |
| Digestible carbohydrates | 64.9 |
| Micronutrients (ash) | 6.2 |
| Energy (kJ/g dry matter) | 17.0 |
| Metabolizable energy (kJ/g dry matter) | 16.4 |
| **Amino acids (% of protein mass)** |  |
| Arginine | 6.3 |
| Histidine | 2.5 |
| Isoleucine | 4.5 |
| Leucine | 7.7 |
| Lysine | 5.4 |
| Methionine | 2.2 |
| Phenylalanine | 4.8 |
| Threonine | 3.7 |
| Tryptophan | 1.1 |
| Valine | 5.0 |
| Alanine | 4.5 |
| Asparagine | 2.6 |
| Aspartic acid | 9.3 |
| Cysteine | 1.6 |
| Glutamine | 2.6 |
| Glutamic acid | 18.0 |
| Glycine | 4.2 |
| Proline | 5.7 |
| Serine | 5.1 |
| Tyrosine | 3.2 |

## References

Banks, I.J., Gibson, W.T. and Cameron, M.M., 2014. Growth rates of black soldier fly larvae fed on fresh human faeces and their implication for improving sanitation. Tropical Medicine and International Health 19: 14–22. 10.1111/tmi.12228

Barragan-Fonseca, K.B., Gort, G., Dicke, M. and van Loon, J.J.A., 2019. Effects of dietary protein and carbohydrate on life-history traits and body protein and fat contents of the black soldier fly Hermetia illucens. Physiological Entomology 44: 148–159. 10.1111/phen.12285

Berggreen, I.E., Schøn, M.L., Nørgaard, J.V. and Jensen, K., 2025. Optimal dietary protein content and essential amino acid limitation in larvae of the black soldier fly (Hermetia illucens). Journal of Insects as Food and Feed 11: 2579–2590. 10.1163/23524588-bja10221

Berner, D., Blanckenhorn, W.U. and Körner, C., 2005. Grasshoppers cope with low host plant quality by compensatory feeding and food selection: N limitation challenged. Oikos 111: 525–533. 10.1111/j.1600-0706.2005.14144.x

Broeckx, L., Frooninckx, L., Slegers, L., Berrens, S., Noyens, I., Goossens, S., Verheyen, G., Wuyts, A. and Van Miert, S., 2021. Growth of black soldier fly larvae reared on organic sidestreams. Sustainability 13: 12953. 10.3390/su132312953

Cammack, J. A. and Tomberlin, J. K., 2017. The impact of diet protein and carbohydrate on select life-history traits of the black soldier fly Hermetia illucens (L.) (Diptera: Stratiomyidae). Insects 8: 56. 10.3390/insects8020056

Capuano, E., Oliviero, T., Fogliano, V. and Pellegrini, N., 2018. Role of the food matrix and digestion on calculation of the actual energy content of food. Nutrition Reviews 76: 274–289. 10.1093/nutrit/nux072

Davis, G.R.F., 1975. Essential dietary amino acids for growth of larvae of the yellow mealworm, Tenebrio molitor L. Journal of Nutrition 105: 1071–1075. 10.1093/jn/105.8.1071

Eggink, K.M., Donoso, I.G. and Dalsgaard, J., 2023. Optimal dietary protein to carbohydrate ratio for black soldier fly (Hermetia illucens) larvae. Journal of Insects as Food and Feed 9: 789–798. 10.3920/JIFF2022.0102

Eriksen, N.T., 2022. Dynamic modelling of feed assimilation, growth, lipid accumulation, and CO_2_ production in black soldier fly larvae. PLoS ONE 17: e0276605. 10.1371/journal.pone.0276605

Fuhrmann, A., Gold, M., Loh, R.K., Chu, C.X., Haberkorn, I., Puniamoorthy, N. and Mathys, A., 2025. Physical properties of food waste influence the efficiency of black soldier fly larvae bioconversion via microbial activity. Journal of Environmental Management 387: 125777. 10.1016/j.jenvman.2025.125777

Gligorescu, A., Toft, S., Hauggaard-Nielsen, H., Axelsen, J.A. and Nielsen, S.A., 2019. Development, growth and metabolic rate of Hermetia illucens larvae. Journal of Applied Entomology 143: 875–881. 10.1111/jen.12653

Gold, M., Tomberlin, J.K., Diener, S., Zurbrügg, C. and Mathys, A., 2018. Decomposition of biowaste macronutrients, microbes, and chemicals in black soldier fly larval treatment: a review. Waste Management 82: 302–318. 10.1016/j.wasman.2018.10.022

Goodrich, H.R., Wood, C.M., Wilson, R.W., Clark, T.D., Last, K.B. and Wang, T., 2024. Specific dynamic action: the energy cost of digestion or growth? Journal of Experimental Biology 227: jeb246722. 10.1242/jeb.246722

Hawkins, A.J.S., 1991. Protein turnover: a functional appraisal. Functional Ecology 5: 222–233. 10.2307/2389260

Jakobsen, K. and Thorbekt, G., 1993. The respiratory quotient in relation to fat deposition in fattening–growing pigs. British Journal of Nutrition 69: 333–343. 10.1079/bjn19930037

Janssen, R.H., Vincken, J.P., Van Den Broek, L.A.M., Fogliano, V. and Lakemond, C.M.M., 2017. Nitrogen-to-protein conversion factors for three edible insects: Tenebrio molitor, Alphitobius diaperinus, and Hermetia illucens. Journal of Agricultural and Food Chemistry 65: 2275–2278. 10.1021/acs.jafc.7b00471

Koethe, M., Taubert, J. and Vervuert, I., 2022. Impact of lysine supplementation on growth and development of Hermetia illucens larvae. Journal of Insects as Food and Feed 8: 35–44. 10.3920/JIFF2020.0059

Laganaro, M., Bahrndorff, S. and Eriksen, N.T., 2021. Growth and metabolic performance of black soldier fly larvae grown on low and high-quality substrates. Waste Management 121: 198–205. 10.1016/j.wasman.2020.12.009

Lee, K.P., 2007. The interactive effects of protein quality and macronutrient imbalance on nutrient balancing in an insect herbivore. Journal of Experimental Biology 210: 3236–3244. 10.1242/jeb.008060

Lemme, A. and Klüber, P., 2024. Rethinking amino acid nutrition of black soldier fly larvae (Hermetia illucens) based on insights from an amino acid reduction trial. Insects 15: 862. 10.3390/insects15110862

Liu, C., Wang, C. and Yao, H., 2019. Comprehensive resource utilization of waste using the black soldier fly (Hermetia illucens (L.)) (diptera: Stratiomyidae). Animals 9: 349. 10.3390/ani9060349

Lomonaco, G., Franco, A., De Smet, J., Scieuzo, C., Salvia, R. and Falabella, P., 2024. Larval frass of Hermetia illucens as organic fertilizer: composition and beneficial effects on different crops. Insects 15: 293. 10.3390/insects15040293

Oonincx, D.G.A.B., Gold, M., Bosch, G., Guillaume, J.B., Rumbos, C.I., El Deen, S.N., Sandrock, C., Bellezza Oddon, S., Athanassiou, C.G., Cambra-López, M., Pascual, J.J., Parodi, A.P., Spranghers, T. and Yakti, W., 2025. Bugbook: nutritional requirements for edible insect rearing. Journal of Insects as Food and Feed. In Press. 10.1163/23524588-bja10226

Oonincx, D.G.A.B., van Broekhoven, S., van Huis, A. and van Loon, J.J.A., 2015. Feed conversion, survival and development, and composition of four insect species on diets composed of food by-products. PLoS ONE 10: e0144601. 10.1371/journal.pone.0144601

Parodi, A., De Boer, I.J.M., Gerrits, W.J.J., Van Loon, J.J.A., Heetkamp, M.J.W., Van Schelt, J., Bolhuis, J.E. and Van Zanten, H.H.E., 2020. Bioconversion efficiencies, greenhouse gas and ammonia emissions during black soldier fly rearing – a mass balance approach. Journal of Cleaner Production 271: 122488. 10.1016/J.JCLEPRO.2020.122488

Pimentel, A.C., Montali, A., Bruno, D. and Tettamanti, G., 2017. Metabolic adjustment of the larval fat body in Hermetia illucens to dietary conditions. Journal of Asia-Pacific Entomology 20: 1307–1313. 10.1016/j.aspen.2017.09.017

Posit Team, 2023. RStudio: Integrated development environment for R (2023.9.1.494). Posit Software (PBC). http://www.posit.co/

Purkayastha, D. and Sarkar, S., 2022. Sustainable waste management using black soldier fly larva: a review. International Journal of Environmental Science and Technology 19: 12701–12726. 10.1007/s13762-021-03524-7

R Core Team, 2023. A language and environment for statistical computing. R Foundation for Statistical Computing. https://www.R-project.org.

Raubenheimer, D. and Simpson, S.J., 1997. Integrative models of nutrient balancing: application to insects and vertebrates. Nutrition Research Reviews 10: 151–179. 10.1079/nrr19970009

Schmidt-Nilsen, K., 1997. Animal physiology: adaption and environment. Cambridge University Press, Cambridge, UK.

Schneider, L., Kisinga, B., Stoehr, N., Cord-Landwehr, S., Schulte-Geldermann, E., Moerschbacher, B.M., Eder, K., Jha, R. and Dusel, G., 2025. Dietary protein levels in isoenergetic diets affect the performance, nutrient utilization and retention of nitrogen and amino acids of Hermetia illucens (L.) (Diptera: Stratiomyidae) larvae. Insects 16: 240. 10.3390/insects16030240

Schøn, M.L., Jensen, K., Schow-Madsen, M., Gold, M., Mathys, A., Berggreen, I.E., Schou, T.M., Nørgaard, J.V. and Overgaard, J., 2025a. Using gas exchange measurements to monitor growth, energy expenditure, and body composition of black soldier fly larvae (Hermetia illucens) on diets varying in protein-to-carbohydrate ratio. bioRxiv. 10.1101/2025.09.08.674872

Schøn, M.L., Mikkelsen, M.V.N., Jensen, K., Poulsen, J.M., Berggreen, I.E., Schou, T.M., Nørgaard, J.V. and Overgaard, J., 2025b. Effect of temperature on growth, metabolism, and gas exchange in Hermetia illucens larvae reared under commercial and laboratory conditions. Journal of Insects as Food and Feed 11: 1059–1074. 10.1163/23524588-00001268

Secor, S.M., 2009. Specific dynamic action: a review of the postprandial metabolic response. Journal of Comparative Physiology B 179: 1–56. 10.1007/s00360-008-0283-7

Siddiqui, S.A., Süfer, Ö., Çalışkan Koç, G., Lutuf, H., Rahayu, T., Castro-Muñoz, R. and Fernando, I., 2024. Enhancing the bioconversion rate and end products of black soldier fly (BSF) treatment – a comprehensive review. Environment, Development and Sustainability 27: 9673–9741. 10.1007/s10668-023-04306-6

Spranghers, T., Moradei, A., Vynckier, K., Boudrez, M., Pinotti, L. and Ottoboni, M., 2025. Amino acid requirements of yellow mealworm and black soldier fly larvae. Journal of Insects as Food and Feed 11: 1047–1058. 10.1163/23524588-00001271

Stryer, L., 1995. Biochemistry. W.H. Freeman. New York, NY.

Talal, S., Cease, A., Farington, R., Medina, H.E., Rojas, J. and Harrison, J., 2021. High carbohydrate diet ingestion increases post-meal lipid synthesis and drives respiratory exchange ratios above 1. Journal of Experimental Biology 224: jeb240010. 10.1242/jeb.240010

Talal, S., Harrison, J.F., Farington, R., Youngblood, J.P., Medina, H.E., Overson, R. and Cease, A.J., 2024. Body mass and growth rates predict protein intake across animals. eLife 12: e88933. 10.7554/eLife.88933

Tomberlin, J.K., Miranda, C., Flint, C., Harris, E. and Wu, G., 2023. Nutrients limit production of insects for food and feed: an emphasis on nutritionally essential amino acids. Animal Frontiers 13: 64–70. 10.1093/af/vfad032

van der Heide, M.E., Stødkilde, L., Nørgaard, J.V. and Studnitz, M., 2021. The potential of locally-sourced European protein sources for organic monogastric production: a review of forage crop extracts, seaweed, starfish, mussel, and insects. Sustainability 13: 2303. 10.3390/su13042303

Van’t Hof, H.M. and Martin, M.M., 1989. The effect of diet water content on energy expenditure by third-instar Manduca sexta larvae (Lepidoptera: Sphingidae). Journal of Insect Physiology 35: 433–436. 10.1016/0022-1910(89)90118-2

Von Bertalanffy, L., 1957. Quantitative laws in metabolism and growth. Quarterly Review of Biology 32: 217–231. 10.1086/401873

Withers, P.C., 2001. Design, calibration and calculation for flow-through respirometry systems. Australian Journal of Zoology 49: 445–461. 10.1071/ZO00057

Wu, G., Bazer, F.W., Dai, Z., Li, D., Wang, J. and Wu, Z., 2014. Amino acid nutrition in animals: protein synthesis and beyond. Annual Review of Animal Biosciences 2: 387–417. 10.1146/annurev-animal-022513-114113

